# Metagenomic association analysis of gut symbiont *Lactobacillus reuteri* without host-specific genome isolation

**DOI:** 10.1101/2020.05.19.101097

**Authors:** Sein Park, Martin Steinegger, Ho-Seong Cho, Jongsik Chun

## Abstract

*Lactobacillus reuteri* is a model symbiont colonizing the guts of vertebrates used for studies on host adaptation of the gut symbiont. Previous studies investigated host-specific phylogenetic and functional properties by isolating its genomic sequence. This dependency on genome isolation is a significant bottleneck. Here we propose a method to study the association between *L. reuteri* and its hosts directly from metagenomic reads without strain isolation by using pan-genomes.

We characterized the host-specificity of *L. reuteri* in metagenomic samples not only in the previously studied organisms (mice and pigs) but additionally in dogs. For each sample, two types of profiles were generated: (1) genome-based strain type abundance profiles and (2) gene composition profiles. Our profiles showed host-association of *L. reuteri* in both phylogenetic and functional aspects without depending on the host-specific genome isolation. We could observe not only the presence of host-specific lineages but also the dominant lineages associated with the different hosts.

Furthermore, we show that metagenome-assembled genomes provide detailed insights into the host-specificity of *L. reuteri*. We could infer evolutionary trajectories of host-associative *L. reuteri* strains in the metagenomic samples by placing the metagenome-assembled genomes into a phylogenetic tree and identify novel host-specific genes which were unannotated in existing pan-genome databases.

Our pan-genomic approach drops the need for time-consuming and expensive host-specific genome isolation while producing consistent results with previous host-association findings in mice and pigs. Additionally, we could predict associations that have not yet been studied in dogs.

## Introduction

*Lactobacillus reuteri* is a Gram-positive bacterial symbiont colonizing the gut in a variety of vertebrate species and is used as a model organism to study the evolutionary process of vertebrate gut symbionts [1, 2]. The evolutionary trajectories of *L. reuteri* were studied through Amplified-Fragment Length Polymorphism (AFLP), Multi-Locus Sequence Analysis (MLSA) and core-genome phylogeny [1, 3, 4]. These studies found genetically distinct subpopulations that highly correlate with their host, indicating a stable host-symbiont relationship. However, they also found some outliers where the strains from unrelated hosts were included in these host-specific clusters, which suggests occasional horizontal transfers between hosts [1, 2]. The transfer from one individual host to another could simultaneously occur in different host populations, resulting in distinct phylogenetic lineages even in the same host species [3].

The adaptation of *L. reuteri* to the respective host resulted in host-specific functional features. For example, comparative genomics analysis on isolates could identify host-specific genes with functions related to transposable elements and biofilm formation [4–6], which was experimentally verified in mice [5–7].

However, the analysis based on the isolated strains might not be able to represent the complete repertoire of host-associated features of *L. reuteri* since isolating and sequencing a single bacterial strain under appropriate culture conditions remains challenging [8]. Currently, there are 187 isolated genomes (April 2020) available in the National Center for Biotechnology Information (NCBI) Genome database [9], but the majority originates from well-studied model species such as rodents, humans, pigs and poultry, limiting our ability to study a wider range of the host adaptation.

In this study, we devised a method (Figure 1) to analyze the association of *L. reuteri* with hosts in metagenomes that overcome the need for the host-specific genome isolation and successfully applied to gut microbiome samples of three mammals: pig, mouse and dog.

**Figure 1.**
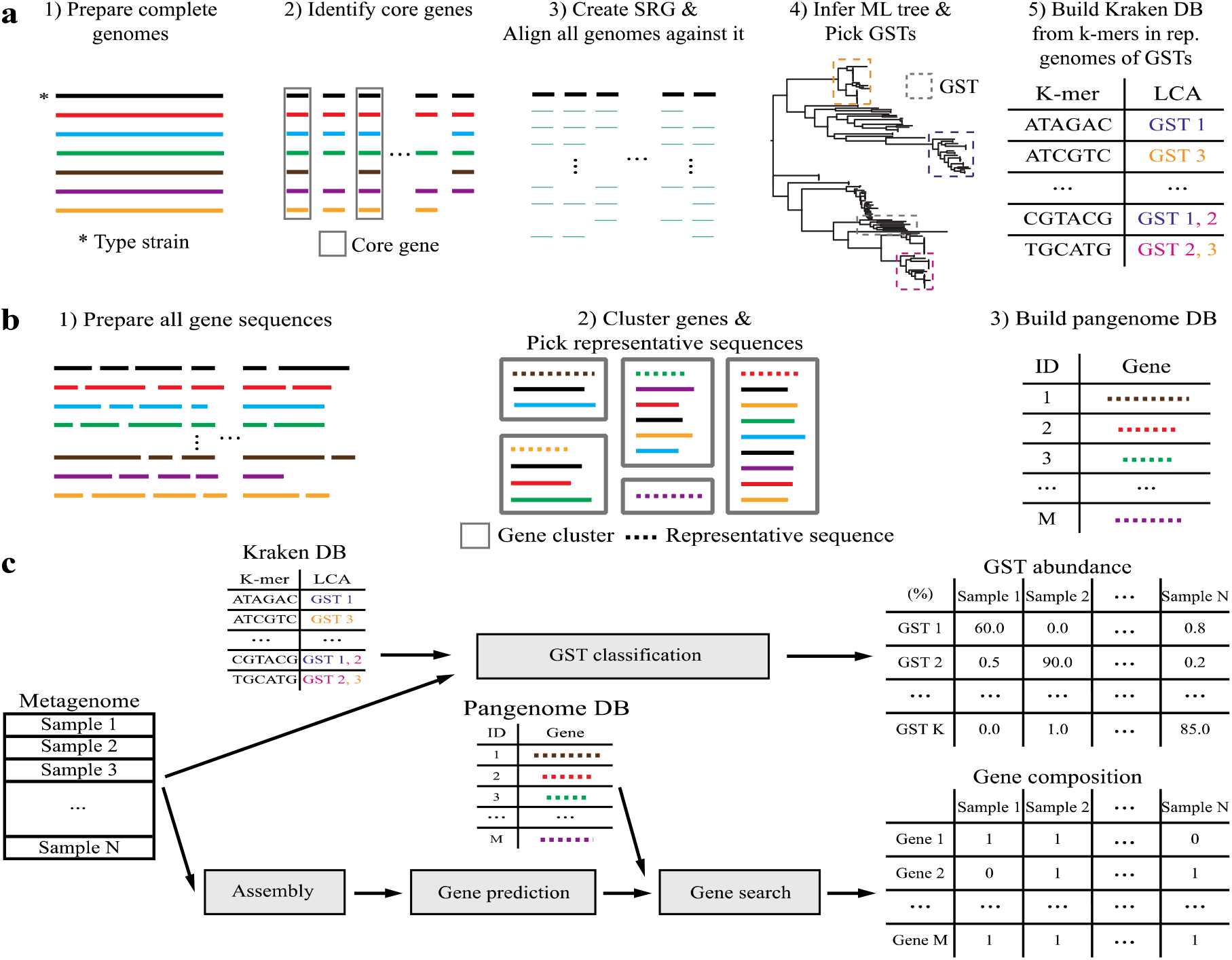
Overview of the analysis. (a) Construction of the Kraken database using the phylogenetic tree. (b) Construction of the pan-genome database. (c) Profiling the metagenome samples based on the reference databases built above.

## Results

### Construction genome-based strain types and gene composition profiles to characterize metagenomic samples

To predict the phylogenetic features of *L. reuteri* in metagenome samples, we profiled the composition of genome-based strain types (GSTs). For this, a species-specific reference genome (SRG), phylogenetic tree and Kraken [10] database were built using complete genomes from the EzBioCloud database [11] (see Additional File 1: Table S1), as described in Figure 1 (a). We created the *L. reuteri* SRG by concatenating 1,158 core genes from eight complete genomes identified by Roary [12], resulting in an 1,097,896 bp long sequence. All available 151 *L. reuteri* (as of April 2020) strains isolated from humans, rodents, pigs, poultry, herbivores (goats, sheep, cows and horses) and food sources (Additional File 1: Table S1) were aligned against the SRG using MUMmer [13]. The resulting multiple sequence alignments were used to infer a maximum likelihood phylogenetic tree using RAxML [14]. We then clustered the tree into 20 GSTs by merging adjacent clades until a maximum all-against-all pairwise distance of 20,000 single nucleotide variations (SNVs) was reached. A reference Kraken database was built by core gene sequences from representative genomes of each GST (Additional File 2: Figure S1).

Moreover, we inferred functional features of *L. reuteri* based on gene composition profiles, which indicate the absence and presence of genes in each sample. As described in Figure 1 (b), a reference pan-genome database was constructed for *L. reuteri* using the coding sequences (CDSs) from 151 genomes. The database is comprised of a collection of 20,014 *L. reuteri*-specific gene clusters, which were obtained by using a 90% DNA similarity and 90% alignment coverage threshold. These clusters include 1,149 core ones found in over 95% of the genomes, 8,926 singletons.

These reference databases were used to characterize the metagenome samples by GST abundance and gene composition (Figure 1 (c)). We profiled each sample by searching its reads against our GST database using Kraken [10] and estimated the abundance using Bracken [15]. The gene composition was profiled through assembly using MEGAHIT [16], genes were predicted using Prodigal [17] and annotated using MMseqs2 [18].

### Evaluation of the profile estimation using synthetic samples

We evaluated the accuracy of GST classification at the read-level and the composition-level using synthetic samples. The synthetic samples were created using InSilicoSeq [19] with three different complexity levels: four low complexity (LC), four middle complexity (MC) and two high complexity (HC) containing either five or ten randomly selected GSTs, or all twenty GSTs, respectively. The precision at the read-level, defined as the proportion of correct assignments in GST and its ancestors to the total number of assignments, was achieved an average precision of 95.68%, 95.16% and 92.39% in the LC, MC and HC samples, respectively (Additional File 1: Table S2). We also measured the accuracy of the composition-level classification by computing the Pearson correlation coefficient between the estimated and true abundance, achieving 0.9937, 0.9879 and 0.9729 on average in the LC, MC and HC samples, respectively (Additional File 3: Figure S2).

Moreover, the accuracy of gene composition profiling was measured using true positive rate (TPR) and F1 score. We simulated metagenomic samples from twelve reference genomes using InSilicoSeq [19] (see Additional File 4: Figure S3 (a) legend) at four different coverage levels (1×, 5×, 10× and 20×) and constructed gene profiles from these samples. We obtained TPRs of ≥ 75% at 5× coverage, and ≥ 90% at 10× and 20× coverage (Additional File 4: Figure S3 (a)), and F1 scores of ≥ 80% at 5× coverage, and ≥ 90% at 10× and 20× coverage (Additional File 4: Figure S3 (b)). In the case of the real metagenomic samples, the coverage was 23.38× on average, with a standard deviation of 22.41 (Additional File 4: Figure S3 (c)).

### Genome-based strain typing to characterize host-associative *L. reuteri* populations in the real metagenomic gut samples

Using the GST abundance profiles created from the real metagenomic gut samples, the samples were found not to contain a mixture of similarly-distributed GSTs but a few abundant GSTs, which showed association with the host origin (Figure 2).

**Figure 2.**
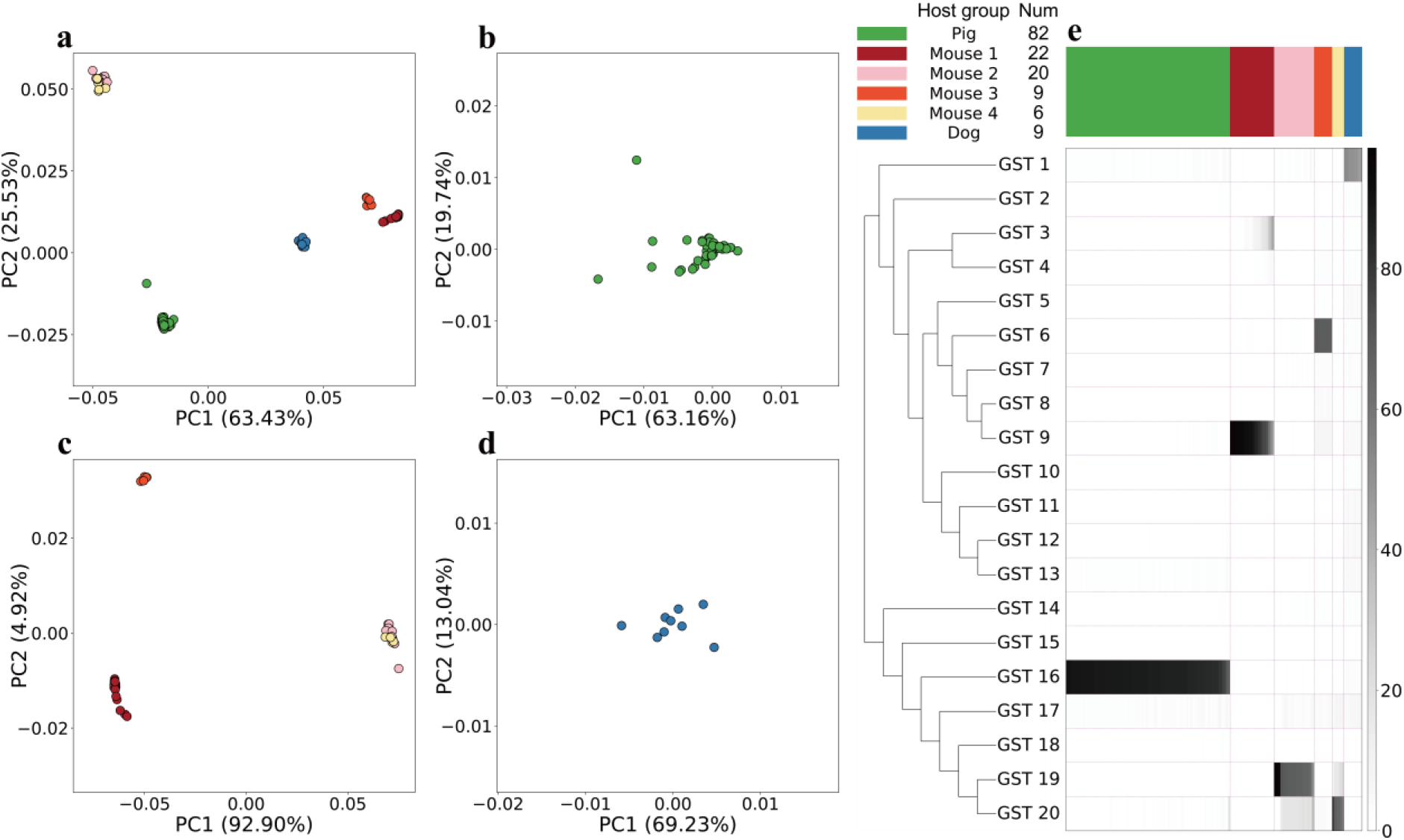
GST abundance profiles. Population structures of *L. reuteri* in (a) all samples, (b) pig samples, (c) mouse samples and (d) dog samples are visualized as Principal Coordinate Analysis (PCoA) plots with weighted UniFrac distance metric. (e) GST abundance profiles are represented as a heatmap, visualizing relative abundance of the GSTs. Phylogenetic relationships between the GSTs are illustrated as a tree on the left (branch lengths are ignored). The host groups are shown in different colors on the top.

Since distinct host-specific lineages could be found from the same host origin, the host species were divided into “host groups” based on the most abundant GST (“dominant GST”) in the metagenomic samples. We identified six host groups: one “Pig” group, four “Mouse” groups and one “Dog” group (Figure 2 (a)-(d)). The dominant GSTs of each host group were GST 16 in “Pig” samples (n = 82), GST 9 in “Mouse 1” samples (n = 22), GST 19 in “Mouse 2” samples (n = 20), GST 6 in “Mouse 3” samples (n = 9), GST 20 in “Mouse 4” samples (n = 6) and GST 1 in “Dog” samples (n = 9) (Figure 2 (e), Additional File 5: Figure S4). Except for the dog samples for which isolated genome sequences were unavailable in the reference database, the host group of the metagenome samples was associated with the isolation sources of the dominant GSTs of the samples (Additional File 2: Figure S1).

These dominant GSTs were consistently found in the placement of 85 medium-to-high quality MAGs into the phylogenetic tree (Figure 3). Unlike the GST profiles, the MAG placements represented the phylogenetic relationship between *L. reuteri* in the samples and reference strains. For example, the MAGs from the pig samples were completely placed into the GST 16 clade (Figure 3 (a)), whereas those from the dog samples formed their own clade outside the reference GST 1 clade (Figure 3 (d)).

**Figure 3.**
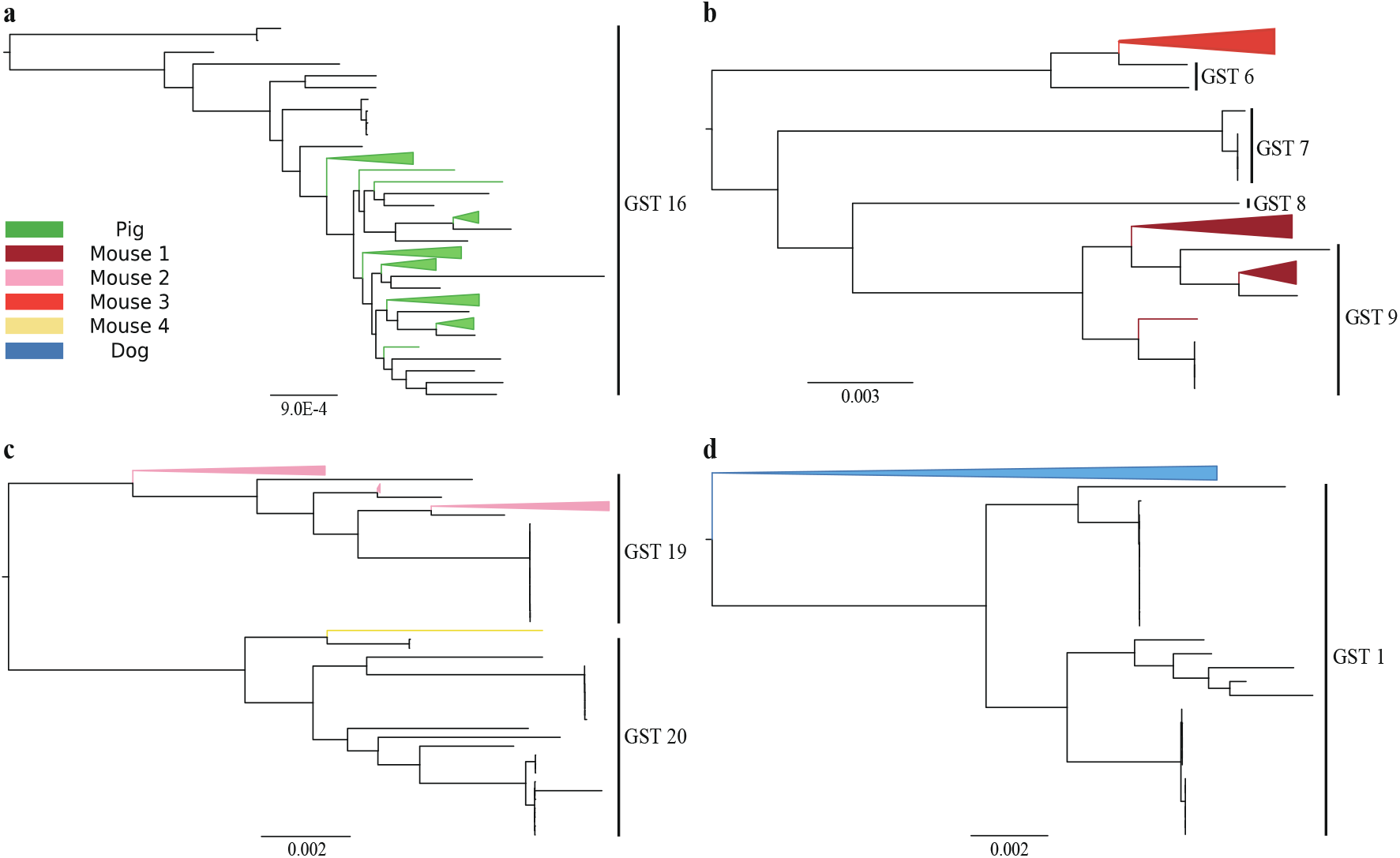
Phylogenetic placement of the MAGs. Phylogenetic trees display the placement of *L. reuteri* MAGs assembled from the reads of the metagenome samples in the (a) “Pig” group, (b) “Mouse 1” and “Mouse 3” groups, (c) “Mouse 2” and “Mouse 4” groups and (d) “Dog” group, respectively.

The relative abundance of the dominant GSTs was different in each host group: the median abundance in “Pig” samples, “Mouse 1” to “Mouse 4” samples and “Dog” samples was 87%, 90%, 68%, 68%, 66% and 50%, respectively (Additional File 5: Figure S4). This abundance indicated that a single dominant GST occupied more than half of the *L. reuteri* population despite the variation in the abundance. However, we additionally found non-dominant GSTs with a relative abundance above 10%. “Mouse 2” and “Mouse 4” samples contained 18% of GST 20 and 17% of GST 19, respectively. Not only the isolation sources of the dominant GSTs but the non-dominant GSTs coincided with the host groups of the samples (Additional File 2: Figure S1).

Furthermore, we checked how distinct each *L. reuteri* population between host groups was by performing a PERMANOVA [20] test. It revealed that the GST abundance of the samples was significantly different from those of other samples included in the different host groups (Additional File 6: Figure S5).

### Functional features of *L. reuteri* associated with host origin

Phylogenetically we could measure the host-association of *L. reuteri* using our GST profiles. Furthermore, we also wanted to see if the host-specificity was reflected in the functional profiles. To investigate this, we picked host-specific genes (HSGs) from our gene composition profiles.

We detected significant differences in *L. reuteri* gene composition between the host groups using the PERMANOVA test (Additional File 7: Figure S6). A set of 4,128 host-specific genes, including 2,172 “Pig” HSGs, 540 “Mouse 1” HSGs, 644 “Mouse 2” HSGs, 354 “Mouse 3” HSGs, 207 “Mouse 4” HSGs and 211 “Dog” HSGs were identified using Fisher’s exact test, and assigned to clusters of orthologous groups (COG) [21] annotation (Figure 4 (a), Additional File 1: Table S3). These HSGs mainly belonged to four functional categories: 1) Replication, recombination and repair, 2) transcription, 3) transport and metabolism of various macromolecules and ions and 4) cell wall/membrane/envelope biogenesis.

**Figure 4.**
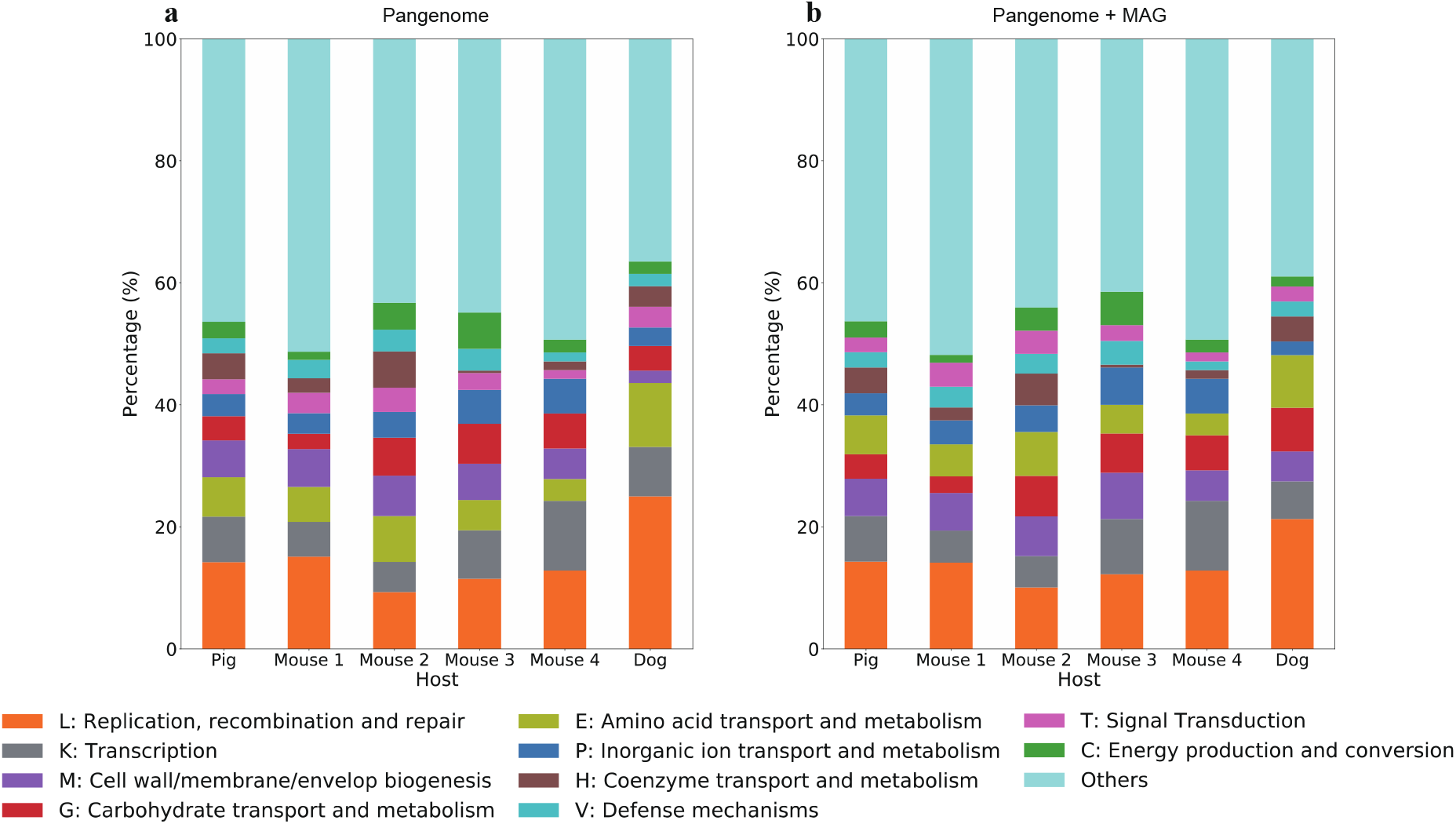
Host-specific functional structures of each host group. The stacked barplots visualize the proportion of the functional categories of host-specific genes. (a) represents the host-specific genes identified from the pangenome database, while (b) represents those identified from the pangenome database and the MAGs.

We hypothesized that many HSGs might not be represented by our reference pan-genome database. Therefore, we predicted 132,255 CDSs from 85 MAGs and searched them against the pan-genome database. On average, the MAGs contained 60.7% of genes were also found in the gene profiles of the corresponding samples; 10.4 % of them were found in the pan-genome database but not in their corresponding gene profiles and 28.9% of them could not be matched to the pan-genome database. These 17,669 unmatched sequences were clustered into 12,648 gene clusters, indicating that 38.72% of gene sequences could not be annotated through searching against the pan-genome database.

By performing Fisher’s exact test, we identified a set of 1,914 HSGs, and 419 of them were newly found in MAGs. These novel HSGs were annotated based on the eggNOG [22] database (Additional File 1: Table S4), and the proportion of COG functional categories of all HSGs was computed, as represented in Figure 4 (b). We compared the functional structures of HSGs based on the pan-genome to those based on both pan-genomes and MAGs, finding that the ten most abundant categories were conserved despite some differences in the ratio (Figure 4). However, if the host-specific reference genome was absent, a relatively high percentage of HSGs were newly identified from the MAGs. About 37% of dog HSGs were exclusively found in the MAGs, while 2% of pig HSGs and 20% of mouse HSGs were only found in the MAGs (Additional File 1: Table S5).

From the detailed functional description in Additional File 1: Table S3 and Table S4, we observed that transposases, integrases and ABC transporters were found to be host-specific in all host groups. However, some gene functions were not identified from all host groups; for example, host-specific urea amidohydrolases were observed only in the “Mouse 1” and “Mouse 2” groups. These host-specific functions, especially those related to mobile elements and biofilm formation, would reflect differences in the host gut environment and the adaptation mechanism of *L. reuteri* to their host, which consistently supports the observation of previous studies with isolated strains [6, 7].

## Discussion

We observed that the placement of the MAGs into the phylogenetic tree provided detailed insights into the phylogenetic relationship between *L. reuteri* in the sample and reference strains. For example, the presence of an independent clade of dog MAGs suggested that the divergence between human-specific and herbivore-specific lineages in reference GST 1 strains was preceded by the divergence of dog-specific lineage. However, the GST profiles based on read-mapping could not clearly explain this evolutionary trajectory since they simply represented the dominance of GST 1 in the dog samples.

Nonetheless, the analysis based on read-mapping has advantages over the MAG placement based one. First, obtaining high-quality MAGs requires a large number of samples [23]. In this study, we obtained 85 medium-to-high quality MAGs from 148 metagenomic samples. Since the quality of MAGs depends on the quality of assembly [23, 24], the phylogenetic placement would be inaccurate if the assembled contigs contain too much noise or have low coverages. Additionally, the MAG often represents multiple strains rather than an individual so that it could cause a bias in the strain-level analysis [23, 24].

The MAGs also provided many novel functional features. A set of 12,648 gene clusters were newly identified, while the pan-genome included 20,014 clusters, meaning that the MAGs contained more than 50% of gene information compared to the pan-genome from isolates. Moreover, about 10% of the HSGs were novel in the MAGs in that 4,128 and 419 HSGs were identified from the pan-genome and MAGs, respectively. In particular, if the host-specific reference genome is unavailable, such as for dogs in this study, a relatively high proportion of novel HSGs would be found only based on MAGs.

## Conclusions

Our approach used metagenomic reads instead of isolated bacterial genomes to analyze the host-symbiont association, allowing us to apply this approach to gut metagenome from any host, such as dogs in this study. We investigated the host-specific population structures and functional features of *L. reuteri* in the metagenomic samples through the reference pan-genome and MAGs. This approach not only demonstrated a consistent association with previously studied hosts, but we were also able to apply it to the host without existing isolated strain from it.

## Methods

### Prioritization of reference genomes

We prepared 151 high-quality reference *L. reuteri* genomes from the EzBioCloud database [11] and sorted them by the level of genome assembly completeness, prioritizing complete genomes, followed by chromosome-level assembled genomes and then others. Those in the same assembly completeness level were re-sorted by their N50 value.

### Phylogenetic analysis and strain typing of *Lactobacillus reuteri*

The maximum likelihood (ML) phylogenetic tree of *L. reuteri* was created through the workflow described by Ha et al. [25]. First, fourteen complete reference genomes were picked, and UBCG core genes [26] were extracted from each genome. We aligned the UBCGs for each complete genome pair, computing the similarity. The genomes of which the median similarity was 100% in all pairs were grouped, and those with the highest priority in each group were selected as representative genomes. Eight resultant representatives were used for *L. reuteri* core-genome identification. We generated a set of core genes by adding only the genes shared among the complete genomes of *L. reuteri* using the Roary v3.12.0 pipeline [12]. A Species-specific Reference Genome (SRG) for *L. reuteri* was then created by concatenating the core gene set. We created a multiple sequence alignment from Single Nucleotide Variations of all 151 genomes aligned against our SRG using MUMmer v3.23 [13]. An ML tree was inferred by RAxML v8.2.8 using the GTRCAT nucleotide model [14].

The genome-based strain types (GSTs) are defined as the largest clades where the number of SNVs were less than 20,000 in all pairs of the clade members, were clustered from *L. reuteri* genomes. Starting from the type strain genome, we merged the sister clades on the ML tree into the GST as long as the maximum number of SNVs between the clade members were below 20,000. The merging step was repeated until everything was clustered, starting from the genome with the highest priority. We selected the genome with the highest priority of each GST as the representative.

Using the kraken-build script in the Kraken software package [10], a database with a k-mer length of 31 was constructed from the core genes of the representative genomes. Since Kraken by default uses the NCBI taxonomy to assign k-mers to a taxonomic level, we provided a custom taxonomy following the topology of our ML tree to map k-mers to the lowest common ancestor (LCA).

### Construction of *L. reuteri* reference pan-genome

To produce an *L. reuteri* specific gene database, we collected all protein-coding sequences (CDSs) from *Lactobacillus* from the EzBioCloud database and clustered them into orthologous groups using Linclust [27] at 90% sequence identity and 90% bi-directional coverage thresholds. Only representative sequences of the clusters that contained *L. reuteri* were added to our pan-genome database and assigned a COG based on the annotation in the EzBioCloud database [11].

### Metagenomic sample collection and sequencing

We collected metagenomic sequence data not only by downloading previously deposited data of pig, mouse and dog gut microbiomes from NCBI SRA database [28–31] but also by directly sequencing 20 pig samples (Additional File 1: Table S6).

Total DNA from fecal samples was isolated using FastDNA Spin Kit for Soil (MP Biomedicals), and its quality was checked using Qubit dsDNA HS Assay Kit (Thermo Fisher Scientific). DNA libraries for whole-genome shotgun sequencing were prepared according to Illumina Truseq Nano protocol, and the 2 × 100 paired-end sequencing was performed using an Illumina NovaSeq 6000 system.

### Metagenomic sequence data processing

Species-level taxonomy profiles were created using a curated core gene-based bacterial database [32]. All samples not containing *L. reuteri* in their profiles were discarded.

Quality filtering and trimming were performed by BBDuk from the BBTools suite [33]. Adapter sequences and low-quality bases from the termini at a quality threshold of Q12 were trimmed from the reads, followed by the removal of the reads with average quality below 10. All reads passing the quality-control filtering were assembled into contigs, using MEGAHIT v1.1.3 [16] with default k-mer sizes and a minimum contig length of 500bp. We used Prodigal v2.6.2 [17] in metagenomic mode to extract all genes from the assembly that longer than 100bp.

### Profiling GST abundance and gene composition in the metagenomic samples

The reads in each metagenomic sample were mapped to the reference Kraken database and results were summarized using kraken-report. We used Bracken [15] to estimate GST abundance from the reports. The gene composition was profiled by searching the predicted genes from the assembled reads against *L. reuteri* pan-genome using MMseqs2 [18] with 90% minimum sequence identity and bi-directional alignment coverage thresholds.

For the real metagenomic samples, we removed those containing less than 20,000 reads classified as *L. reuteri* or less than 500 predicted gene hits. The remaining samples were clustered by their host group, those including the same dominant GSTs. The host group and its samples were filtered out if it did not include more than five samples.

### Metagenome binning and picking *L. reuteri* MAGs

MetaBAT2 v2.14 [34] with the option “--minContig 1500” was used for contig binning. The coverage information was provided through Bowtie2 [35], which mapped the reads into the contigs and SAMtools [36], which converted the mapping results into BAM format. CheckM v1.1.1 [37] was used for assessing the completeness and contamination of each genome bin with UBCGs [26] as a marker gene set, selecting medium-quality genomes as having completeness of ≥ 50% and contamination of < 10%, and high-quality genomes as having completeness of > 90% and contamination of < 5% [38]. We calculated Average Nucleotide Identity (ANI) values between the type strain genome of *L. reuteri* and metagenome-assembled genomes (MAGs) using the OrthoANIu tool [39], assigning the MAGs with ANI value > 95% to *L. reuteri* [40].

### Validation with simulated metagenome data

We generated synthetic datasets for three complexity levels from the *L. reuteri* reference genome sequences using InSilicoSeq v1.3.5 [19]: (1) Four low-complexity (LC) samples, (2) four middle-complexity (MC) samples and (3) two high-complexity (HC) samples. The LC, MC and HC samples respectively contain 10 million reads with five randomly chosen GSTs, 50 million reads with ten randomly chosen GSTs and 50 million reads with all twenty GSTs. The genome abundance was log-normally distributed in half of the samples and exponentially distributed in the remaining ones.

The read-level accuracy to identify *L. reuteri* GSTs in the metagenomic samples was assessed by a precision metric. We computed the precision by dividing the number of assignments to the correct GST and its ancestors by the total number of assignments. The composition-level accuracy of the GST profiles was evaluated based on Pearson’s correlation coefficient. We calculated the coefficient between true and estimated GST abundance.

Furthermore, the true positive rates (TPRs) and F1 scores were used for evaluating the accuracy of profiling *L. reuteri* gene composition. A total of twelve complete isolate genomes were simulated into synthetic metagenome reads at four coverage levels: 1×, 5×, 10× and 20×. These synthetic reads were assembled using MEGAHIT [16], and CDSs were predicted from contigs ≥ 500bp using Prodigal [17], as described above. Gene composition profiles for each simulated sample were constructed by searching CDSs ≥ 100bp against the pan-genome database using MMseqs2 [18] with 90% minimum sequence identity and bi-directional alignment coverage thresholds. To compare the gene profiles to the true gene composition of isolated genomes, we defined TPRs and F1 scores as:

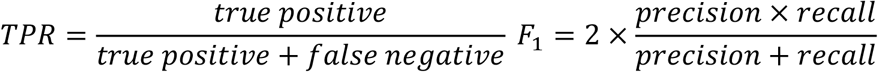

### Phylogenetic placement and novel gene identification of MAGs

SNVs were detected from the medium and high-quality *L. reuteri* MAGs using MUMmer [13], merged with the SNVs from 151 *L. reuteri* genomes into MSA. We placed the MAGs into the phylogenetic tree of *L. reuteri* using RAxML with the “-f v” option [14]. Genes of the MAGs, predicted by Prodigal [17], were searched against the *L. reuteri* pan-genome database using MMseqs2 [18] and compared to the gene composition profiles of the corresponding metagenome samples to identify novel genes of MAGs. The unmatched genes were clustered using Linclust [27] with 90% of minimum sequence identity and bidirectional alignment coverage thresholds and annotated with eggNOG [22] assignments using eggNOG-mapper v2.0.1 [41].

### Statistical analysis

Permutational multivariate analysis of variance (PERMANOVA) [20] with 9,999 permutations was used for determining if the GST abundance and gene composition of the samples from one host group were significantly different from those from another. We used the weighted UniFrac distance [42] for GST profiles and the Jaccard distance for gene composition profiles as distance metrics.

For each host and gene, a contingency table representing the difference between the presence and absence of the genes in the gene composition profiles of the samples from the host was created and used for Fisher’s exact test to identify host-specific genes (HSGs). We determined the gene to be host-specific if the p-value was below < 0.01.

## List of abbreviations

GST: Genome-based strain type
HSG: Host-specific gene

## Declarations

### Ethics approval and consent to participate

Not applicable.

### Consent for publication

Not applicable.

## Availability of data and materials

Raw metagenomic data are available from the SRA database (see Additional File 1: Table S6). Codes for picking GSTs, building Kraken and pan-genome databases and profiling metagenomic data are available in https://github.com/psi103706/Lreuteri_strain_analysis.

## Authors’ contributions

SP implemented the software and performed the research, SP, MS, JC designed the research, SP, MS, JC drafted the manuscript, HC generated the pig metagenomic data. All authors read and approved the final manuscript.

## Acknowledgements

The authors would like to thank Milot Mirdita for his helpful comments and proofreading the manuscript.

## Supplementary Information

**Additional File 1. Supplementary Tables.** This file contains Table S1 to Table S6.

**Additional File 2. Figure S1. Maximum likelihood phylogenetic tree of *L. reuteri* strains with GSTs and isolation sources.**

The outgroups, *Lactobacillus vaginalis* ATCC 49540, *Lactobacillus panis* DSM 6035 and *Lactobacillus frumenti* DSM 13145, are not shown in the figure. The branches are colored by the isolation source of the strains.

**Additional File 3. Figure S2. Correlation between the estimated and true GST abundance of the synthetic samples.**

**Additional File 4. Figure S3. The accuracy of gene search with respect to the coverage depth.**

The plots report (a) true positive rate and (b) F1 score at the coverage of 1×, 5×, 10× and 20×, respectively. Synthetic data simulated from isolate genomes of twelve *L. reuteri* strains were used for the evaluation. (c) represents the distribution of the coverage level of the real metagenomic samples.

**Additional File 5. Figure S4. GST abundance of the host groups.**

The boxplots represent the GST abundance of (a) “Pig” group, (b) “Mouse 1” group, (c) “Mouse 2” group, (d) “Mouse 3” group, (e) “Mouse 4” group and (f) “Dog” group, respectively.

**Additional File 6. Figure S5. Comparison of GST abundance using weighted UniFrac distance.**

The plots represent the distance from (a) “Pig” group, (b) “Mouse 1” group, (c) “Mouse 2” group, (d) “Mouse 3” group, (e) “Mouse 4” group and (f) “Dog” group, respectively. P-values of the PERMANOVA test were shown above each box in the plots (* : < 0.01, ** : < 0.001, ***: < 0.0001)

**Additional File 7. Figure S6. Comparison of gene composition using Jaccard distance.**

The plots represent the distance from (a) “Pig” group, (b) “Mouse 1” group, (c) “Mouse 2” group, (d) “Mouse 3” group, (e) “Mouse 4” group and (f) “Dog” group, respectively. P-values of the PERMANOVA test were shown above each box in the plots (* : < 0.01, ** : < 0.001, ***: < 0.0001)

