## Supplementary figures and images for "Metagenomic association analysis of gut symbiont *Lactobacillus reuteri* without host-specific genome isolation"

### Figure S1

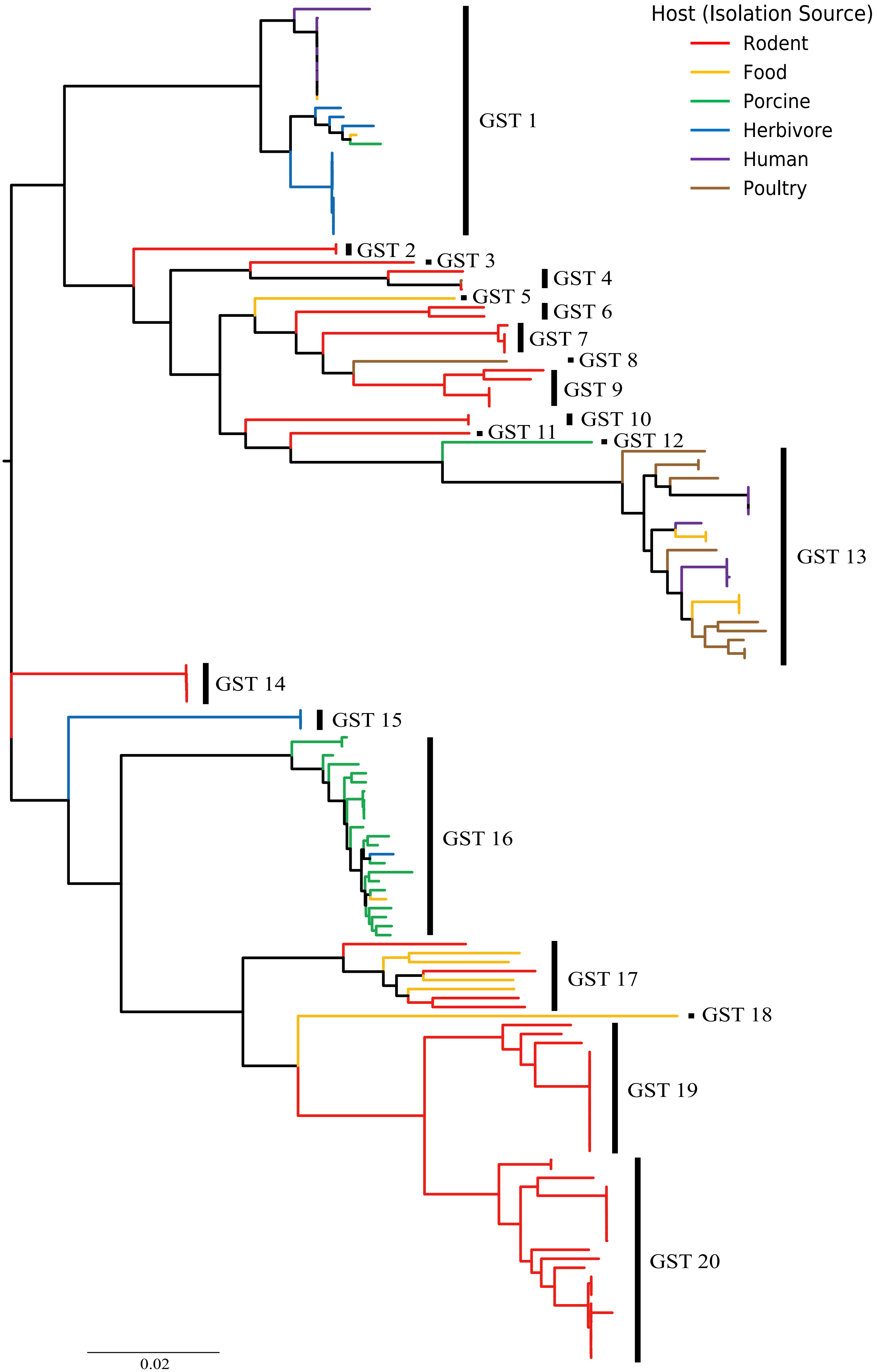

### Figure S2

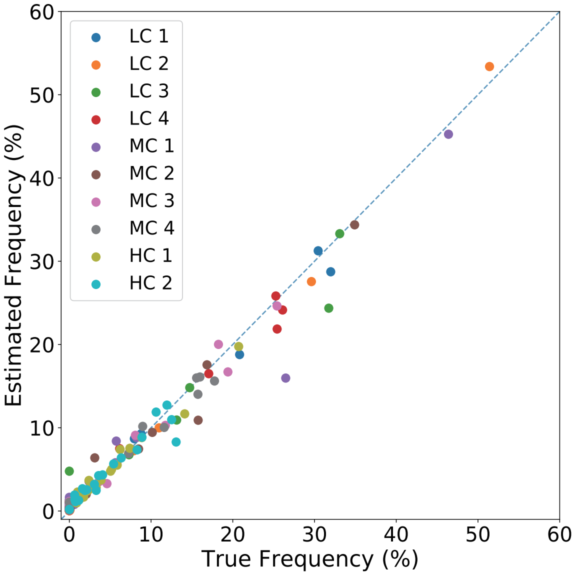

### Figure S3

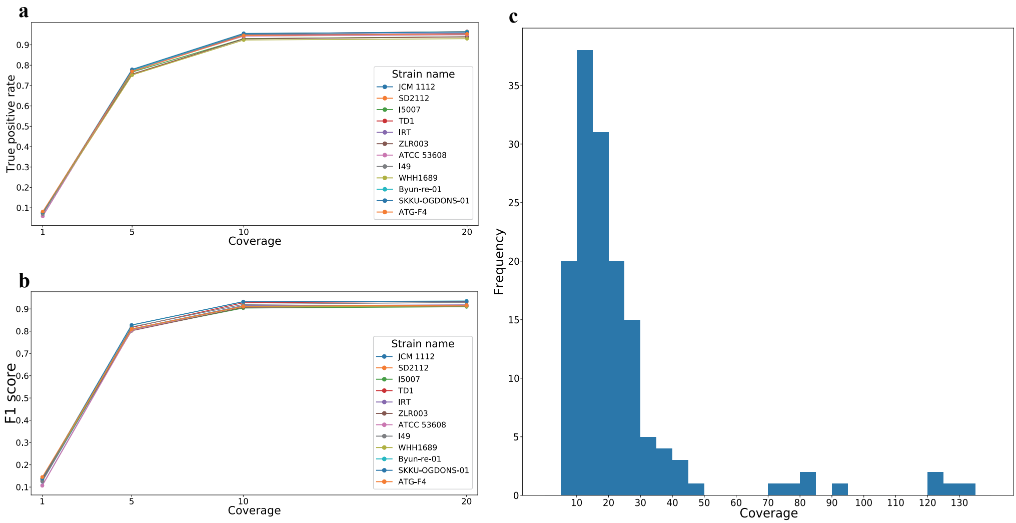

### Figure S4

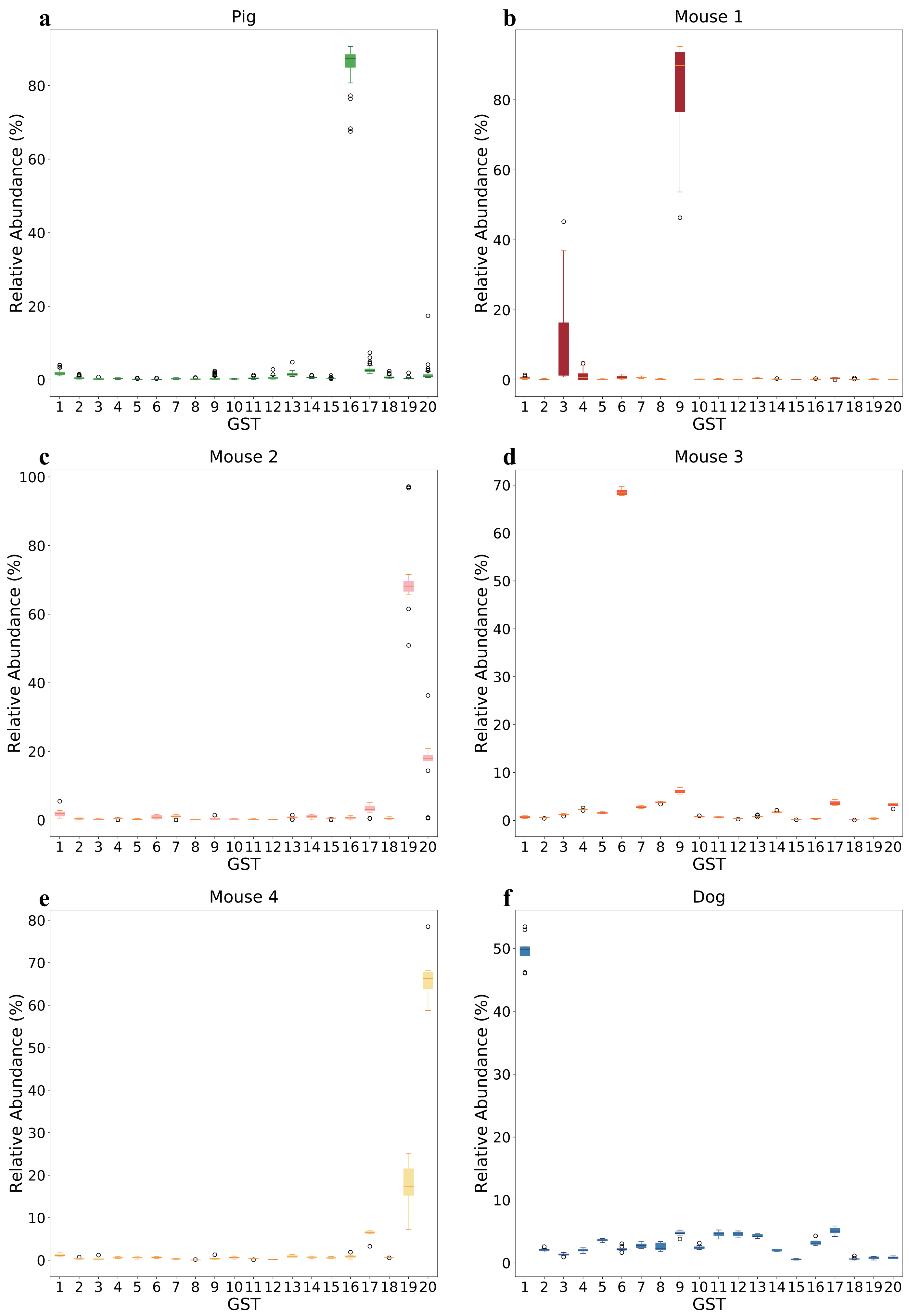

### Figure S5

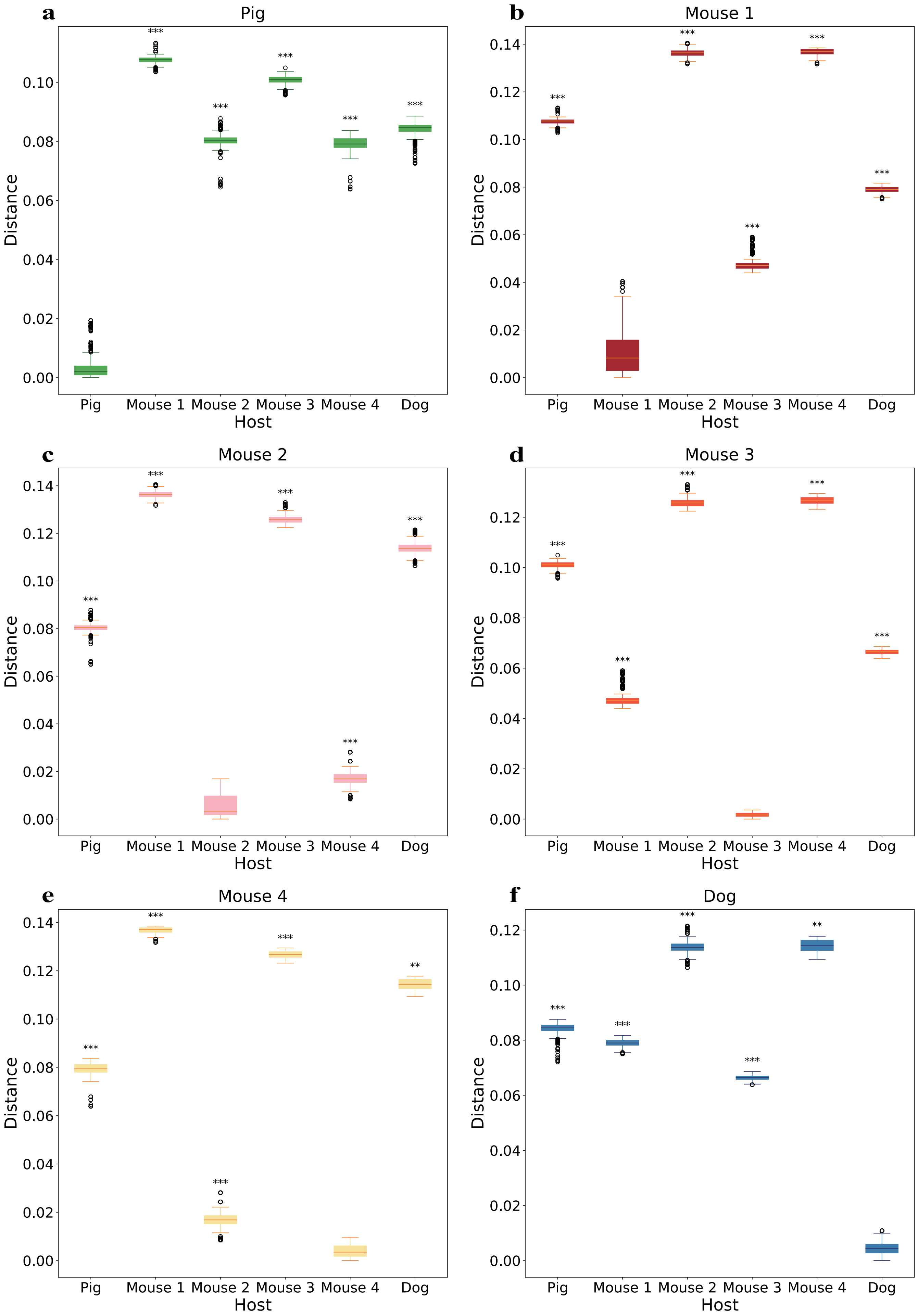

### Figure S6

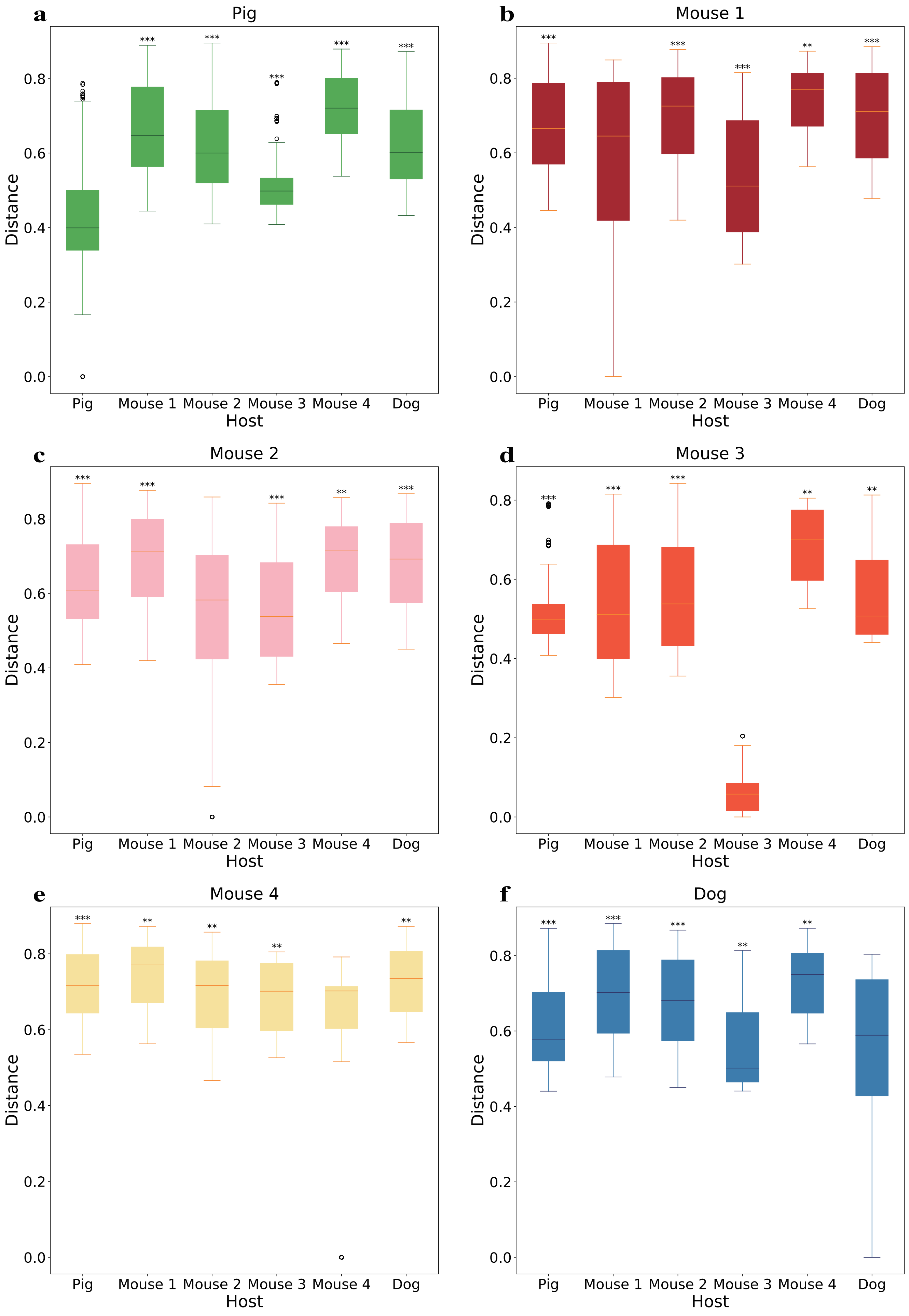
